# Strongly truncated *Dnaaf4* plays a conserved role in *Drosophila* ciliary dynein assembly as part of an R2TP-like co-chaperone complex with *Dnaaf6*

**DOI:** 10.1101/2022.05.12.491607

**Authors:** Jennifer Lennon, Petra zur Lage, Alex von Kriegsheim, Andrew P. Jarman

## Abstract

Axonemal dynein motors are large multi-subunit complexes that drive ciliary movement. Cytoplasmic assembly of these motor complexes involves several co-chaperones, some of which are related to the R2TP co-chaperone complex. Mutations of these genes in humans cause the motile ciliopathy, Primary Ciliary Dyskinesia (PCD), but their different roles are not completely known. Two such dynein (axonemal) assembly factors (DNAAFs) that are thought to function together in an R2TP-like complex are DNAAF4 (DYX1C1) and DNAAF6 (PIH1D3). Here we investigate the *Drosophila* homologues, *CG14921*/*Dnaaf4* and *CG5048*/*Dnaaf6*. Surprisingly, *Drosophila* Dnaaf4 is truncated such that it completely lacks a TPR domain, which in human DNAAF4 is likely required to recruit HSP90. Despite this, we provide evidence that *Drosophila* Dnaaf4 and Dnaaf6 proteins can associate in an R2TP-like complex that has a conserved role in dynein assembly. Both are specifically expressed and required during the development of the two *Drosophila* cell types with motile cilia: mechanosensory chordotonal neurons and sperm. Flies that lack either gene are viable but with impaired chordotonal neuron function and lack motile sperm. We provide molecular evidence that *Dnaaf4* and *Dnaaf6* are required for assembly of outer dynein arms (ODAs) and a subset of inner dynein arms (IDAs).

## INTRODUCTION

Ciliary motility is driven by a highly conserved family of axonemal dynein motors, which are large multi-subunit complexes (King, 2016). Those that comprise the Outer Dynein Arms (ODA) are the main drivers of motility, whereas those of the Inner Dynein Arms (IDA) modulate ciliary movement. During ciliogenesis, the assembly of the motors into the cilium or flagellum is highly regulated. After subunit synthesis, complex assembly occurs within the cytoplasm (known as pre-assembly) prior to transport and docking within the cilium (Fok et al., 1994; Fowkes and Mitchell, 1998). This pre-assembly is facilitated by a series of regulators called dynein pre-assembly factors (DNAAFs) (King, 2016). Many of these factors were originally identified as causative genes of human Primary Ciliary Dyskinesia (PCD), but they are highly conserved among eukaryotes that have motile ciliated cells (Omran et al., 2008). This conservation was recently shown to be true for *Drosophila melanogaster*, which has an almost full complement of homologous genes for the axonemal dynein complexes and for dynein assembly factors (zur Lage et al., 2019). In the case of *Drosophila*, ciliary motility is confined to the sensory cilium of mechanosensory neurons (chordotonal neurons) and the sperm flagellum. Flies with dysfunctional dyneins are therefore deaf, uncoordinated and have immotile sperm, which makes the fly a convenient model for analysis of motile ciliogenesis (Diggle et al., 2014; Moore et al., 2013; zur Lage et al., 2018; zur Lage et al., 2021).

The specific functions of DNAAFs are beginning to be unravelled, and in many cases they are thought to function as co-chaperones that regulate HSP70/90 to facilitate correct folding of the dynein heavy chains as well as subunit assembly (Fabczak and Osinka, 2019). Chaperones are important for many cellular functions including the assembly of large multi-subunit complexes like axonemal dynein motors. For several DNAAFs, such a function is strongly indicated by DNAAF sequence relationships with a known HSP90 co-chaperone, the R2TP complex (Maurizy et al., 2018). This co-chaperone was discovered in *S. cerevisiae* as facilitating RNA polymerase II assembly (Zhao et al., 2005). In humans, R2TP comprises the ATPases Ruvbl1 and Ruvbl2, a TPR (tetratricopeptide repeat) protein RPAP3, and a Pih domain protein PIH1D1 (Table 1). R2TP facilitates the assembly/stabilisation of several multi-subunit complexes, including RNA polymerase II, PIKKs (Kakihara and Houry, 2012). Much is known of the structural features of R2TP: for RPAP3, the TPR domains directly recruit HSP70 and HSP90 while the RPAP3_C domain binds to PIH1D1. For PIH1D1, the PIH domain recruits client proteins, while the CS domain binds to RPAP3 (Kakihara and Houry, 2012; Maurizy et al., 2018). Similarly, in a proteomic profiling of *Chlamydomonas* mutants, *mot48* (PIH1D1) *pf13* (DNAAF2) and *twi* (DNAAF6) have overlapping but distinct roles in assembly of dynein complex subsets (Yamamoto et al., 2010; Yamamoto et al., 2020).

**Table 1.** Genes referred to in this study.

| *Drosophila* | Flybase ID | Human* | *C. reinhardtii* | Danio rerio |
| --- | --- | --- | --- | --- |
| *pontin (pont)* | FBgn0040078 | *RUVBL1* | *CrRuvBL1* | *ruvbl1* |
| *reptin (rept)* | FBgn0040075 | *RUVBL2* | *CrRuvBL2* | *ruvbl2* |
| *spaghetti/Rpap3* | FBgn0015544 | *RPAP3* | *Cr02.g084900* | *rpap3* |
| *Spag1* | FBgn0039463 | *SPAG1* | *Spag1* | *spag1a/b* |
| *CG14921/Dnaaf4* | FBgn0032345 | *DNAAF4 (DYX1C1)* | *pf23/DYX1C1* | *dnaaf4* |
| *Pih1D1* | FBgn0032455 | *PIH1D1* | *mot48?* | *pih1d1* |
| *CG4022/Pih1D2* | FBgn0035986 | *PIH1D2* | *n/a* | *pih1d2* |
| *CG5048/Dnaaf6* | FBgn0036437 | *DNAAF6 (PIH1D3)* | *twi(?)* | *twister* |
| *Nop17l* | FBgn0033224 | *DNAAF2/KTU* | *pf13* | *ktu* |

\*Following nomenclature recommendations in Braschi et al. (Braschi et al., 2022).

There is evidence that mutation of Ruvbl1/2 also causes ciliary dynein defects (Li et al., 2017; Zhao et al., 2013). While this may partly be due to involvement of R2TP in dynein pre-assembly as has been demonstrated in zebrafish and *Drosophila* (Yamaguchi et al., 2018; zur Lage et al., 2018), it is thought that Ruvbl1/2 may also function with DNAAFs to form ‘R2TP-like’ complexes specifically required for dynein assembly (Fig. 1A) (Olcese et al., 2017; Pal et al., 2014; Vaughan, 2014). Among the DNAAFs, SPAG1 has both TPR and RPAP3_C domains, while DNAAF4 (DYX1C1) has TPR and CS domains. Similarly, the CS and PIH domains of PIH1D1 are also present in several other PIH proteins: PIH1D2, DNAAF2 (KTU), and DNAAF6 (PIH1D3) (Dong et al., 2014). There is biochemical evidence that SPAG1 complexes with PIH1D2 and DNAAF2 (Maurizy et al., 2018; Smith et al., 2022). Different isoforms of DNAAF4 complex with DNAAF2 and DNAAF6 (Maurizy et al., 2018; Olcese et al., 2017; Paff et al., 2017; Tarkar et al., 2013). Whether these putative complexes function *in vivo* and their precise role during dynein assembly are not fully established, but they may be required for different steps in the process or for the assembly of different dynein subtypes. For the PIH proteins, the possibility of different roles during dynein assembly has been raised by experiments in zebrafish and *Chlamydomonas* (Yamaguchi et al., 2018; Yamamoto et al., 2010; Yamamoto et al., 2020). In zebrafish, *pih1d1, pih1d2* and *ktu* and *twister* (DNAAF6 homologue) have overlapping functions in the assembly of ODAs and IDA subsets based on analyses of mutant spermatozoa (Yamaguchi et al., 2018).

**Figure 1.**
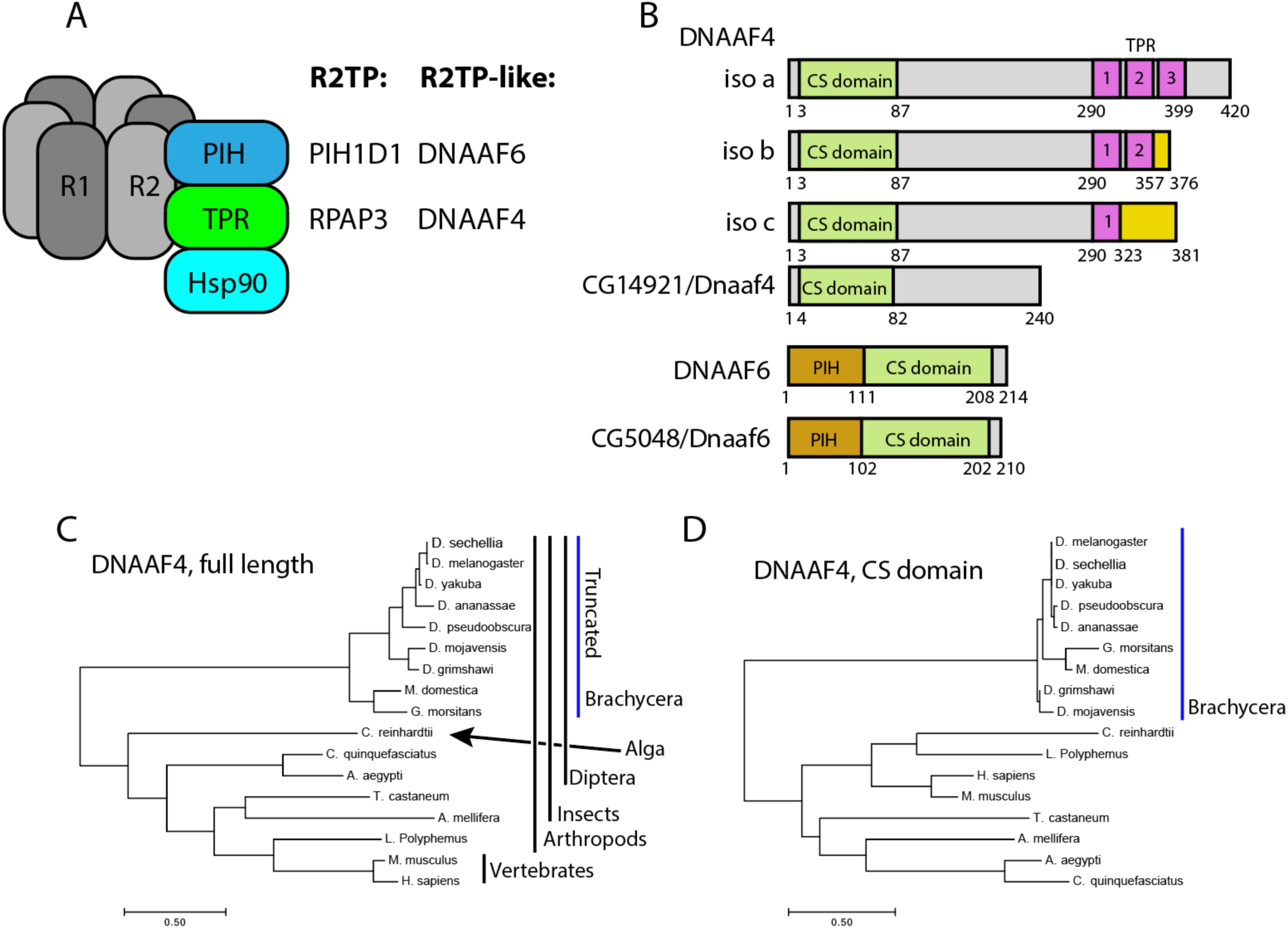
*DXrosophila* and mammalian Dnaaf4/Dnaaf6 proteins. (A) Schematic showing the composition of R2TP and putative DNAAF4/6-containing R2TP-like complexes. (B) Schematic showing the protein domains of human DNAAF4, DNAAF6 and their *Drosophila* orthologues. Human isoforms and protein structures are based on Maurizy et al. (2018). (C) Phylogenetic tree of DNAAF4 sequences from selected species including vertebrates, arthropods and the unicellular green alga, *Chlamydomonas reinhardtii* (established ciliary motility model organism). Higher dipterans (Brachycera) form a distinct group that correlates with gene truncation (blue bar). (D) When comparing CS domains alone, the tree structure remains similar, with Brachycera distinct from other taxa. Organisms included in this tree: *Drosophila sechellia, D. melanogaster, D. yakuba, D. ananassae, D. pseudoobscura, D. mojavensis, D. grimshawi, Musca domestica, Glossina morsitans, Culex quinquefasciatus, Aedes aegypti, Tribolium castaneum, Apis mellifera, Chlamydomonas reinhardtii, Limulus polyphemus, Mus musculus* and *Homo sapiens*.

Of the TPR-containing DNAAFs, *DNAAF4* is a cause of PCD in humans, with motile cilia showing reduction in subsets of ODAs and IDAs (Tarkar et al., 2013). In *Chlamydomonas* the DNAAF4 homologue also shows a partial reduction in ODAs and some IDAs (Yamamoto et al., 2017). In addition to this ciliary motility role, DNAAF4 was originally identified (as DYX1C1) as being affected by a chromosomal translocation associated with susceptibility to developmental dyslexia (Taipale et al., 2003), and subsequently a role for this gene in cortical neuron migration was proposed (Wang et al., 2006). Neither function has an obvious direct link to ciliary motility, suggesting that DNAAF4 may have wider roles beyond dynein pre-assembly. Similarly, SPAG1 may have roles in addition to dynein pre-assembly: R2SP complexes with PIH1D2 were characterised in cells that lack motile cilia (Chagot et al., 2019; Maurizy et al., 2018), and a constitutively expressed isoform exists (Horani et al., 2018).

Thus, the roles of TPR- and PIH-domain containing DNAAFs in assembling subsets of dynein complexes remain to be fully disentangled, as do the identities of the R2TP-like complexes that function in vivo. Moreover, the question of functions for TPR subunits (and by extension the complexes) beyond dynein assembly also remains open.

We have previously shown that *Drosophila* has homologues of *SPAG1* and *DNAAF4* (zur Lage et al., 2019) (Table 1), and that *Drosophila* Spag1 is required for dynein assembly and is able to form a complex with Ruvbl1/2 and Pih1d1 (zur Lage et al., 2018). However, the predicted Dnaaf4 protein is truncated such that it lacks any TPR domain, bringing into question its ability to function in a co-chaperone complex. *Drosophila* has homologues of all the PIH proteins (zur Lage et al., 2019). Most *Drosophila* PIH genes appear widely expressed, but *Dnaaf6* expression appears to be restricted to motile cilia cells. Here we characterise the function of *Drosophila Dnaaf4* and *Dnaaf6* as potential R2TP-like partners. Despite the truncation of Dnaaf4, we show that Dnaaf4 and Dnaaf6 proteins can form an R2TP-like complex, and that each is required for assembly of ODAs and a subset of IDAs. Moreover, there is no indication of functions other than dynein assembly.

## MATERIALS AND METHODS

### Fly stocks

Fly stocks were maintained on standard media at 25°C. The following UAS RNAi stocks were obtained from the Vienna *Drosophila* Resource Center (Dietzl et al., 2007): KK60100 (genetic background stock used as negative control) KK111069 (*Dnaaf4*), KK108561 (*Dnaaf6*) and KK100470 (*Spag1*). The following were obtained from the Bloomington Drosophila Stock Centre: Or-R as wild-type control (#2376), UAS-*Dcr2* (#24644), *w*^*1118*^ *y*^*1*^ *M{vas-Cas9} ZH-2A/FM7c* (#51323), *y*^*1*^ *w* P{y*^*t7*.*7*^*=nos-phiC31\int*.*NLS}X; P{y*^*t7*.*7*^*=CaryP}attP40* (#79604) and *w*; P{UASp-Venus*.*GAP43}7* (#30897). Dnali1-mVenus, Dnal1-mVenus are described in Xiang et al. (Xiang et al., 2022). Flies with UAS-int attp40 landing site were obtained from the Cambridge Microinjection facility. The *sca-Gal4* line used for sensory neuron knockdown was a gift from M. Mlodzik (Baker et al., 1996) and was used in conjunction with UAS-*Dcr2*. For male germline knockdown, *w; Tft/CyO; Bam-Gal4*-VP16 was a gift from Helen White-Cooper.

### Sequence analyses

For detecting orthology, DIOPT was used (Hu et al., 2011). For phylogenetic analysis, protein sequences were obtained from BLAST, Uniprot (Bateman et al., 2021) and Flybase (Larkin et al., 2021). Sequences were aligned using CLUSTALW/MUSCLE within MEGA7 (Kumar et al., 2016). Tree analysis was conducted using the Maximum Likelihood method within MEGA7.

### In situ hybridisation on whole-mount embryos

Primers were designed to give a probe of around 420-bp with the reverse primer containing the T7 RNA polymerase promoter at its 5’ end (all primers are in Supplementary Table S1). DNA was amplified from genomic DNA by PCR and then DIG-labelled RNA generated (DIG RNA Labelling Mix, Roche Cat. No.11277073910) using T7 RNA polymerase (Roche Cat. No. 10881767001). RNA in situ hybridisation was carried out according to zur Lage et al. (2019). In the case of RNA in situ/antibody staining double labelling, antibody staining was carried out after the ISH had been developed. Images were taken on an Olympus AX70 upright microscope with DIC optics.

### Immunofluorescence

Immunohistochemistry on embryos and pupal antenna was described in zur Lage et al. (zur Lage et al., 2018). *Drosophila* testis fixing and staining was carried out according to Sitaram et al. (Sitaram et al., 2014). The following primary antibodies were used: goat anti-GFP antibody (1:500, ab6673), rabbit anti-GFP antibody (1:500, Life Technologies, A11122), mouse anti-Futsch antibody (1:200, Developmental Studies Hybridoma Bank, 22C10), mouse anti-pan polyglycylated tubulin (1:100, Merck, MABS276), rabbit anti-Sas-4 (1:350, gift from Jordan Raff) and rabbit anti-Dnah5 antibody (1:2000, (zur Lage et al., 2021)). The following secondary antibodies were used: goat anti-Rabbit antibody (1:500, Alexa Fluor 488, Life Technologies, A11008) and goat anti-Mouse antibody (1:500, Alexa Fluor 568, Life Technologies, A11019), donkey anti-goat antibody (1:500, Alexa Fluor 488, Life Technologies, A11055), donkey anti-mouse antibody (1:500, Alexa Fluor 568, Life Technologies, A10037), and donkey anti-rabbit antibody (1:500, Alexa Fluor 647, Life Technologies, A31573). Phalloidin was used 1:2000 (Life Technologies, A12380). DNA in adult testes was stained with To-Pro-3 (1:1000, Life Technologies, T3605) or DAPI (14.3mM, Life Technologies) solution in the dark for 15min. After several washes, the samples were mounted on slides with 85% glycerol and 2.5% propyl gallate (Sigma, P3130). Images were captured using a Zeiss LSM-5 PASCAL/Axioskop 2 and a Leica TCS SP8 confocal microscope and processed with Fiji.

### mVenus fusion gene construction

mVenus fusion genes were constructed for Dnaaf4 and Dnaaf6 by amplifying gene segments from genomic DNA and cloning into pDONR221 using the BP clonase II from Gateway technology (Thermo Fisher Scientific). The segment included introns, 5’ UTR, TSS, and additional upstream flanking DNA of approximately 1 kb, but lacked the stop codon. The insert was subsequently transferred to the destination vector pBID-GV (modified from pBID-UASC-GV vector (Wang et al. 2012) where the UASC had been deleted) with the help of LR clonase II (Gateway technology, Thermo Fisher Scientific). This put the ORF in-frame with the mVenus coding sequence. Transformant fly lines were generated by microinjection into syncytial blastoderm embryos of the attP40 landing site line.

### *Dnaaf4* and *Dnaaf6* CRISPR/Cas9 mutant construction

The CRISPR/Cas9 mutant lines were designed by substituting the coding regions of the gene with the mini-white gene. CRISPR primers were designed using the flyCRISPR OptimalTarget finder programme. The cloning was performed according to Vieillard et al. (Vieillard et al., 2016) and injection into the Cas9 line was carried out by the *Drosophila* Microinjection Services (Department of Genetics, Cambridge, UK).

### Fertility, hearing and climbing assays

These assays were carried out as described in zur Lage et al. (2021). In the fertility assay, individual males were crossed to pairs of virgin OrR females and resulting progeny counted. For climbing assays, 2-5 day-old adult females were tested in batches of 15. For the larval hearing assay, batches of 5 third instar larvae on an agar plate placed on a speaker were tested for response to a 1000-Hz tone. *n* for each genotype = 5 batches of 5 larvae, each exposed to 3 tones 30 s apart. For visual analysis of spermatogenesis, testes were dissected, mounted in PBS, and then observed immediately by DIC optics.

### Protein expression analysis of testes by MS

Knockdown males were generated by crossing UAS-RNAi males from *Dnaaf4, Spag1* and the KK control line to *Bam*-Gal4 at 25°C. 1-3 days post-eclosion male progeny were dissected in ice-cold PBS and 30 pairs of testes with four replicates per genotype were snap-frozen in liquid nitrogen before subsequently being processed and analysed for label-free mass-spectrometry as described in zur Lage et al. (2018). The mass spectrometry proteomics data have been deposited in the ProteomeXchange Consortium via the PRIDE (Perez-Riverol et al., 2019) partner repository with the dataset identifier PXD033608.

### Transmission electron microscopy (TEM)

Adult heads were cut off and the proboscis was removed to facilitate infiltration of the solution. The head were rinsed in 0.1 M phosphate buffer before fixing overnight at 4°C in freshly made 2.5% glutaraldehyde, 2% paraformaldehyde in 0.1M phosphate buffer (pH 7.4) solution. Subsequently the samples were rinsed four times and then washed three times for 20min in 0.1M phosphate buffer at room temperature. Further processing for TEM, post-fixing and imaging was carried by Tracey Davey at the Electron Microscopy Research Services, Newcastle University Medical School, using a Philips CM100 CompuStage (FEI) microscope and an AMT CCD camera.

### Transfection and coIP of S2 cells

RNA was prepared from *Drosophila* antennae or testes and mouse testes with the RNeasy Mini kit (Qiagen 74106). cDNA was synthesised, the open reading frames were PCR amplified and initially cloned into the pDONR221 plasmid using the BP clonase II of the Gateway system (Life Technology) before transferring the fragments using the LR clonase II to the C-terminal site of the destination plasmids pAWH (3xHA epitopes) and pAWF (3x FLAG epitopes) of the *Drosophila* Gateway Vector collection (Carnegie Institution for Science). Primers for synthesis are listed in Table S1. The truncated mouse Dyx1c1DTPR protein contains the first 227 amino acids of the wildtype 420 amino acid protein, therefore omitting the whole of the C-terminal TPR domain and replacing it with a stop codon. Transfection into S2 cells was performed according to the X-TREME GENE HP DNA transfection reagent (Merck) protocol. After 48-72 h cells were harvested and coIP was carried out according to the FLAG Immunoprecipitation kit (Sigma-Aldrich). Samples were run on pre-cast gels (Bio-Rad) followed by Western blotting. The blots were then probed with mouse anti-FlagM2 (1:1,000; F1804; Sigma-Aldrich) and rabbit anti-HA (1:4,000; ab9110; Abcam) antibodies, followed by Li-COR secondary antibodies (IR Dye 680RD and IR Dye 800CW), before protein detection on a Li-COR Odyssey scanner using Image Studio v5.2 software.

### GFP trap affinity purification and mass spectrometry

150 pairs of testes in 3 replicates were dissected in ice-cold PBS for Dnaaf4-mVenus and control line UAS-GAP43-mVenus x Bam-Gal4). The samples were snap-frozen in liquid nitrogen. Lysis buffer (Tris-HCl pH7.5 50mM, NaCl 100mM, Glycerol 10%, EDTA 5mM, sodium deoxycholate 0.5%, Complete Mini protease inhibitor) was added to samples before they were homogenised on ice for 2 minutes. Samples were subsequently rotated, incubated in a lysis buffer for 30 minutes at 4°c, and then centrifuged, before being processed and analysed as described in zur Lage et al. (2018) with following alterations: the data was acquired using a Fusion Lumos mass spectrometer (Thermo Fisher) that was operated in an OT-IT configuration. 1-s cycle time, 120k resolution in the orbitrap for MS and rapid scanning MS/MS in the ion-trap. Collision energy was set to 30.

## RESULTS

### *Drosophila* has orthologues of DNAAF4 and DNAAF6, but the former is strongly truncated thereby lacking a TPR domain

Of the PIH genes in *Drosophila*, the orthology prediction tool DIOPT (Hu et al., 2011) identifies the orthologue of *DNAAF6* as *CG5048* (hereafter named *Dnaaf6*). Predicted *Drosophila* Dnaaf6 protein retains PIH and CS domains, and has 45% similarity and 30% identity with the human protein (Fig. 1B). For *DNAAF4*, DIOPT identifies the gene *CG14921* as the *Drosophila* orthologue. However, the encoded protein of this gene (named *Dnaaf4*) is severely truncated relative to the human protein such that it lacks the C-terminal TPR domain (Fig. 1B). Despite this, DIOPT predicts clear orthology with human DNAAF4 for the remaining protein, with 40% similarity and 25% identity. Moreover, the region of alignment is not limited to the CS domain. Phylogenetic analysis indicates that this truncation occurred during dipteran evolution, as the truncation is shared by other higher dipterans (Brachycera) but not lower dipterans or other insects (Fig. 1C). Interestingly, the DNAAF4 sequences of brachyceran flies form a distinct group in a phylogenetic tree, even if just the CS domains are compared (Fig. 1D). This suggests significant sequence divergence occurred in these truncated *Dnaaf4* genes compared with the archetypal full-length genes present from single celled algae to vertebrates.

Human DNAAF4 binds to HSP90 (Tarkar et al., 2013) and this is predicted to occur via its TPR domain (Haslbeck et al., 2013). The loss of this domain in *Drosophila* Dnaaf4 may therefore be expected to have profound consequences for the conservation of *Drosophila Dnaaf4* function as an R2TP-like chaperone in dynein assembly. Below, this is explored by examining expression, protein interactions and gene function.

### *Drosophila Dnaaf4* and *Dnaaf6* are expressed exclusively in differentiating motile ciliated cells

Transcription of both *Dnaaf4* and *Dnaaf6* is highly specific to tissues with motile ciliated cells. Examination of FlyAtlas 2 transcriptome data (Krause et al., 2022) indicates that *Dnaaf4* is expressed specifically in adult testis. In addition, *Dnaaf4* is 5.2-fold enriched in the transcriptome of developing embryonic chordotonal cells (zur Lage et al., 2019). *Dnaaf6* is also very highly expressed in testis, and found to be enriched in chordotonal cells (55.4-fold). RNA *in situ* hybridisation confirms that embryonic expression of each gene is confined to differentiating chordotonal neurons (Fig. 2A,C). In *Dnaaf4* this expression becomes restricted to a subset of lch5 neurons late in differentiation (Fig. 2B). Expression of *Dnaaf6* was abolished in embryos homozygous for a mutation in *fd3F*, which encodes a transcription factor that regulates motile ciliary genes (Newton et al., 2012) (Fig. 2D).

Expression was confirmed in flies with mVenus fusion transgenes, each including about 1-kb of upstream flanking sequence to drive expression under endogenous regulation (Fig. 2F). In each reporter, there are predicted binding sites for the cilia-associated transcription factors fd3F and Rfx. For both *Dnaaf4* and *Dnaaf6*, fusion protein was detected in embryonic chordotonal neurons (Fig. 2G,H), the differentiating chordotonal neurons of Johnston’s organ (JO) in the pupal antenna (Fig. 2I,J), and also in developing spermatocytes (Fig. 2K,L). The fusion protein was located in the cytoplasm of these cells, consistent with a dynein pre-assembly role.

**Figure 2.**
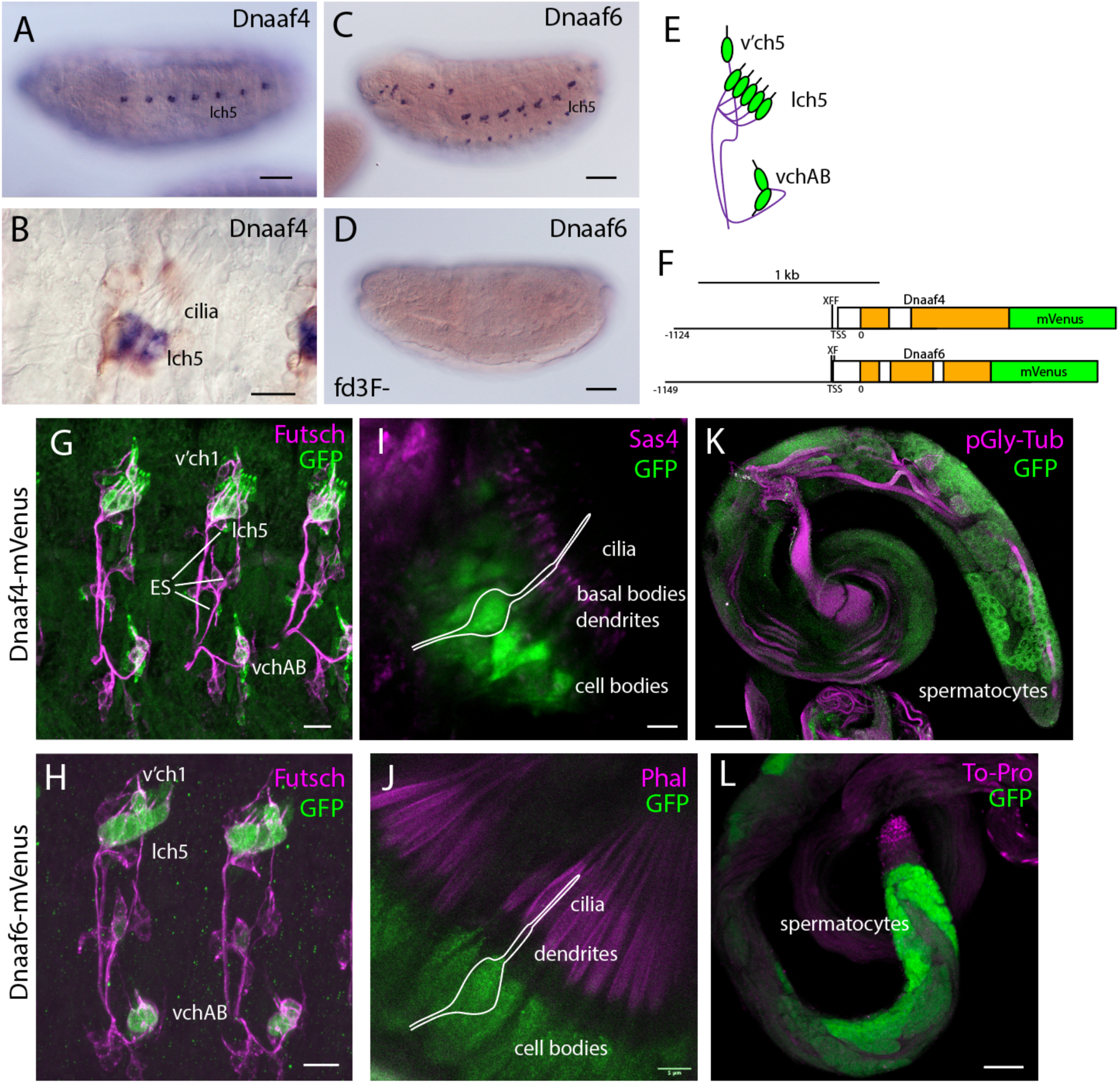
Dnaaf4 and Dnaaf6 are both expressed in *Drosophila* motile cilia cells. (A–D) RNA *in situ* hybridisation conducted on late-stage whole-mount embryos. (A) *Dnaaf4* probe, *Dnaaf4* is expressed specifically in the chordotonal neurons. (B) Higher magnification indicates that this expression becomes restricted at a late stage to a subset of chordotonal neurons (lch5). Here the embryo has been counterstained with antibodies against the sensory neuron marker, Futsch (brown) (C) *Dnaaf6* shows expression in developing chordotonal neurons. (D) In an embryo homozygous mutant for *fd3F*, expression of *Dnaaf6* is abolished. (E) Schematic of the arrangement of chordotonal neurons in embryonic abdominal segments. (F) Schematic illustrating mVenus fusion transgenes. Each includes 5’ flanking DNA containing potential binding sites for the transcription factors fd3f (F) and Rfx (X) (Dnaaf4: C<u>TGTTCAC</u>TTG, <u>GTTCAC</u>TTGCAGC; Dnaaf6: ACTAA<u>ATAAACA</u>A, <u>GTTGCC</u>AGGAAA). (G–L) Expression of Dnaaf4-mVenus detected by anti-GFP antibodies. (G,H) Late embryos counterstained with anti-Futsch (magenta) show expression of both fusion genes in chordotonal neurons. In the case of Dnaaf4-mVenus, some expression is observed in some external sensory (ES) neurons. As this is not observed for the mRNA, it is likely an artefact of the expression construct. (I,J) In pupal antennae, both fusion genes are expressed in the cell bodies of chordotonal neurons that form Johnston’s Organ. A schematic of approximate neuronal location is shown. The counterstain (magenta) is the basal body marker Sas4 (I) or phalloidin (J), which marks the actin basket (scolopale) that surrounds the cilia. (K,L) In adult testes, both fusion genes are expressed in differentiating germline cells (spermatocytes and spermatids). Counterstains (magenta) are polyglycylated tubulin (K) or To-Pro (L). Scale bars are: (A,C,D,K,L) 50 μm (B,G,H) 10 μm (I,J) 5 μm. Number of samples imaged: (G) *n* = 7 (I) *n* = 9 (K) *n* = 8.

In conclusion, despite the truncated nature of Dnaaf4, both proteins are expressed exclusively in motile ciliated cells, consistent with a conserved function in motile ciliogenesis.

### *Drosophila* Dnaaf4 and Dnaaf6 can associate in an R2TP-like complex

Protein interactions were explored by heterologous expression of tagged proteins in S2 cultured cells. Firstly, coimmunoprecipitation confirmed that mouse Dnaaf4 and Dnaaf6 can participate in an R2TP-like complex that also includes Hsp90 (Fig. 3A,B). Similarly, *Drosophila* Dnaaf4 is able to interact with *Drosophila* Dnaaf6 directly, although this interaction appears to be weaker than that between the equivalent mouse proteins (Fig. 3C). Each *Drosophila* protein is also able to complex with Reptin/Pontin (Fig. 3D). Given this association, we asked whether a truncated version of mouse Dnaaf4 retains binding potential. However, this version (mDnaaf4TPR) showed very poor ability to bind to mouse Dnaaf6 (Fig. 3E).

**Figure 3.**
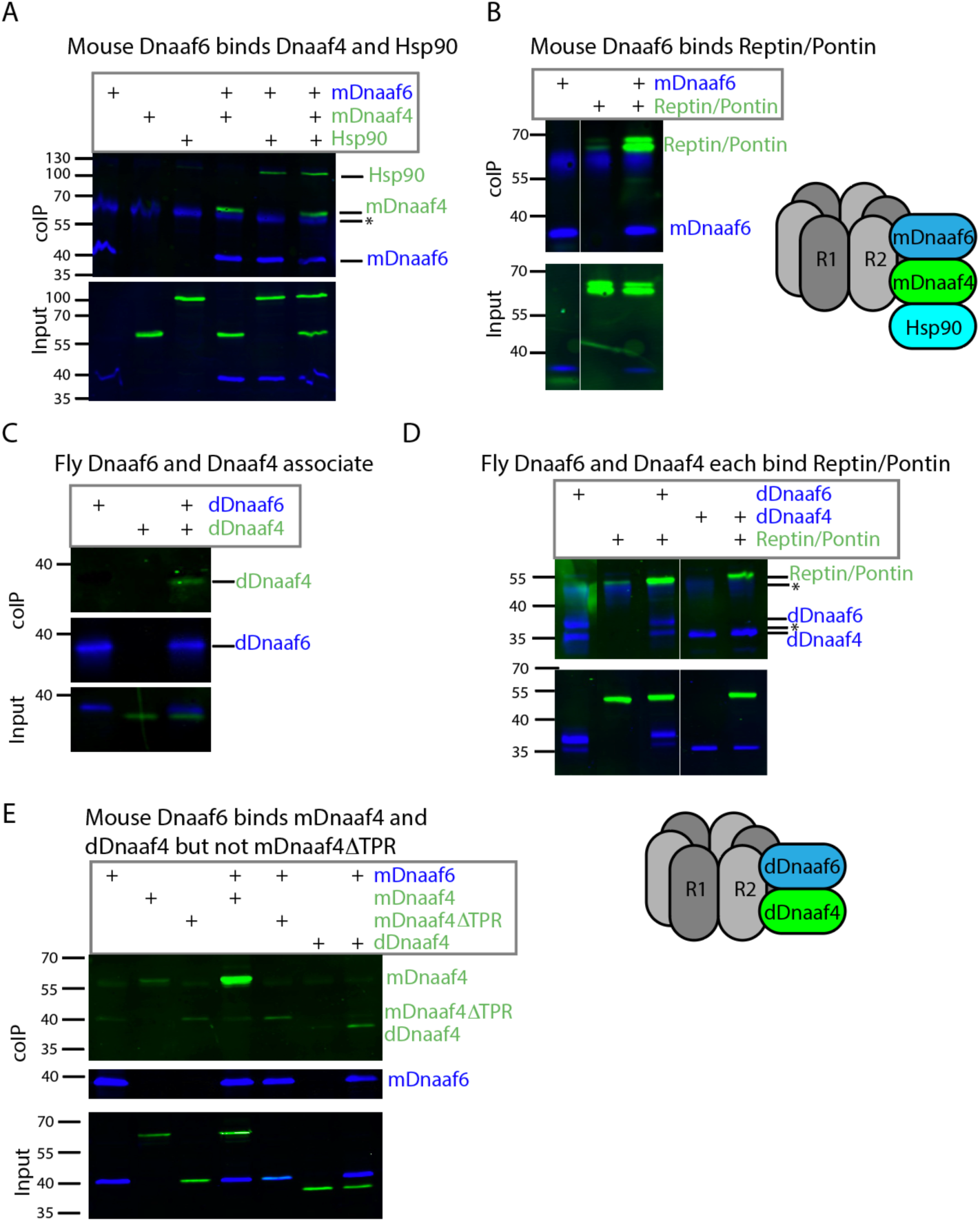
*Drosophila* and mouse Dnaaf4/Dnaaf6 complexes. Coimmunoprecipitations of tagged proteins expressed in S2 cells. In each case, the bait protein is FLAG-tagged (blue) and the prey protein is HA-tagged (green). ‘Input’ represents Western blot of whole cell extracts with bait/prey simultaneously detected (anti-FLAG + anti-HA). ‘coIP’ represents FLAG-mediated coIP followed by simultaneous detection of FLAG- and HA-tagged proteins on Western blot. *indicates non-specific bands. (A) Mouse FLAG-Dnaaf6 protein associates with HA-Dnaaf4 and HA-Hsp90. (B) Mouse FLAG-Dnaaf6 protein binds *Drosophila* HA-Reptin/HA-Pontin. (C) *Drosophila* FLAG-Dnaaf6 and HA-Dnaaf4 associate. (D) *Drosophila* FLAG-Dnaaf6 and FLAG-Dnaaf4 are each capable of binding HA-Reptin/HA-Pontin. (E) Mouse FLAG-Dnaaf6 binds both mouse HA-Dnaaf4 and *Drosophila* HA-Dnaaf4, but is unable to bind the mouse Dnaaf4 protein with TPR domain deleted (HA-Dnaaf4DTPR).

Given the lack of TPR domain in *Drosophila* Dnaaf4, and consequent inability to recruit Hsp90, we searched for protein partners that may provide TPR functionality. A GFP-trap affinity purification was carried out on testes expressing the Dnaaf4-mVenus fusion protein. The associated proteins included Pontin (Fig. 4A). However, of the other associated proteins identified, none appeared to have TPR domains or other features that would help clarify Dnaaf4 function. Filtering the data for proteins associated with motile cilia (zur Lage et al., 2021), we found two proteins of interest to be associated but at a P value that is below the threshold for significance (Fig. 4B). Heatr2 is a known dynein assembly factor (Diggle et al., 2014), while *CG13901* is the *Drosophila* orthologue of mouse *Dpcd*, a gene previously linked to ciliary motility and that associates with R2TP (Dafinger et al., 2018).

**Figure 4.**
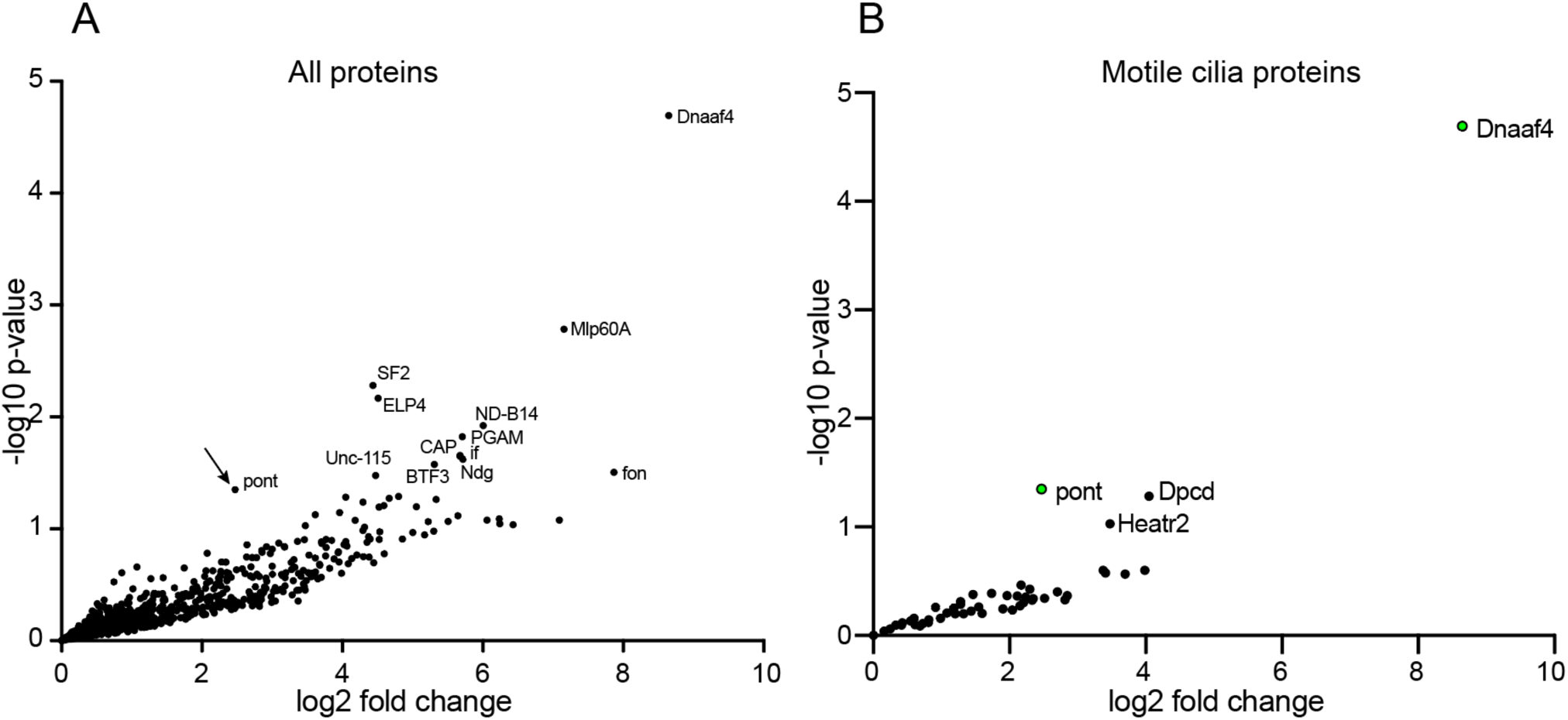
Proteins preferentially associated with Dnaaf4 in *Drosophila* testes. Volcano plots of proteins detected by MS after affinity purification of Dnaaf4-mVenus, shown as relative abundance (fold change) compared with proteins associated with unrelated control protein (GAP43-mVenus). (A) All proteins, with those above threshold significance (-log10(p-value)>1.3) labelled. Pontin of R2TP is significantly associated (arrow). (B) The same dataset filtered to extract proteins associated with motile cilia (zur Lage et al., 2021). Pontin is the only associated protein to reach statistical significance. However, two other proteins of interest are just below significance threshold: Dpcd and Heatr2. Significance was determined using the Empirical Bayes method. *n =* 150 pairs of testes per replicate; 3 replicates per genotype.

### *Dnaaf4* and *Dnaaf6* are required for motile ciliated cell function

To determine the functions of *Dnaaf4* and *Dnaaf6*, we initially examined the effects of knockdown using genetically supplied RNA interference. Knockdown of each gene in the male germline (using *BamGal4* driver) resulted in males that produced significantly fewer progeny than controls (Fig. 5A). A climbing assay was used to test the proprioceptive ability and coordination of adult flies. Knockdown of *Dnaaf4* in sensory neurons (UAS-*Dcr2, scaGal4*, UAS-*Dnaaf4* RNAi^KK111069^) resulted in a significant reduction in climbing ability, consistent with defective chordotonal neuron function (Fig. 6A). Similar reduction was seen for *Dnaaf6* (UAS-*Dcr2, scaGal4*, UAS-*Dnaaf6* RNAi^KK108561^) (Fig. 6B).

**Figure 5.**
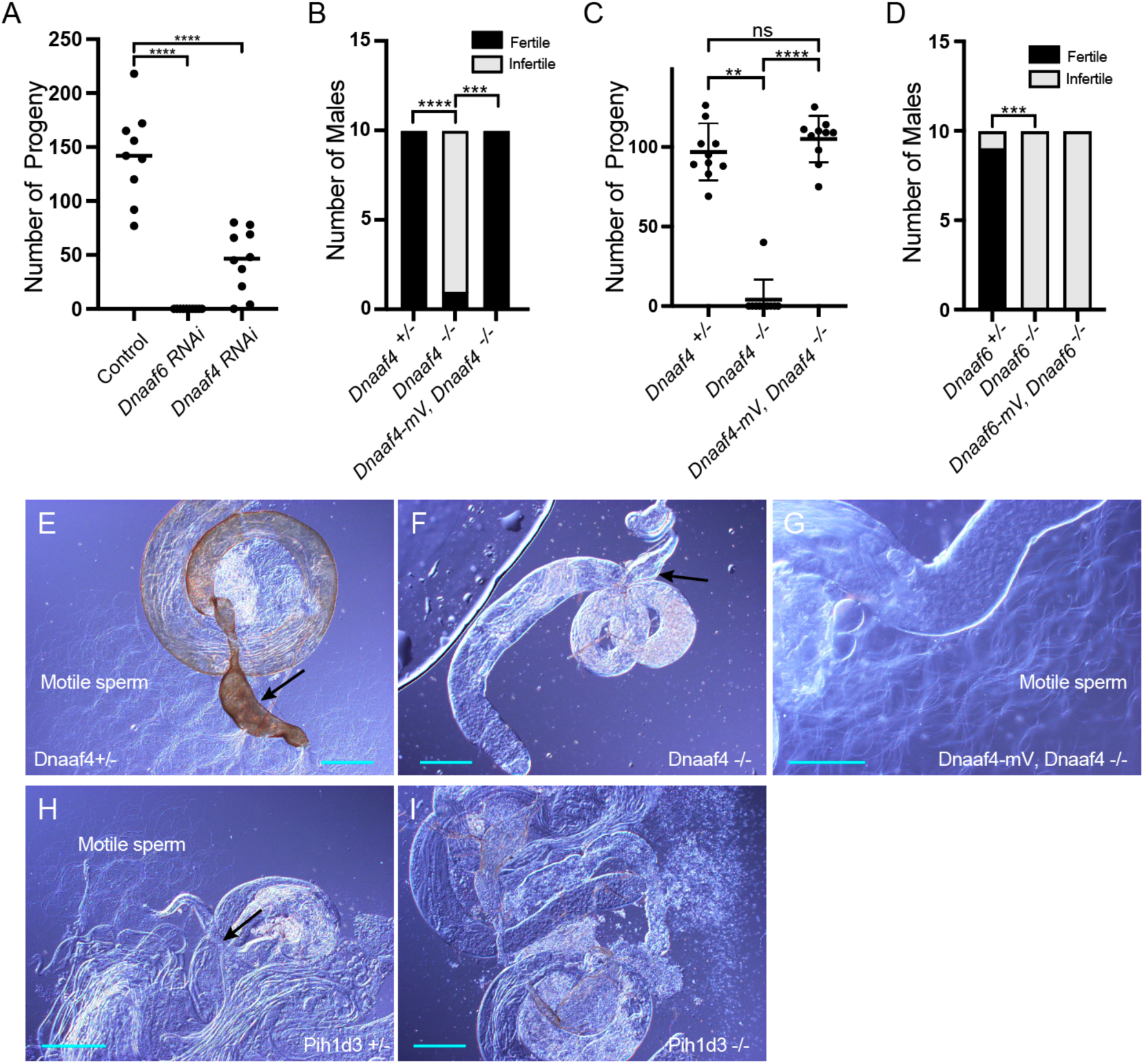
Knockdown and Null mutants of *Dnaaf4* and *Dnaaf6* are male infertile. (A) *Dnaaf4* and *Dnaaf6* RNAi knockdown males (*BamGal4*) produce fewer progeny than control males. Progeny from individual males and median progeny value are shown. Knockdown of either gene significantly reduces progeny per male (P<0.0001, One-way ANOVA followed by Sidak’s Test for multiple comparisons). (B,C) Fertility of *Dnaaf4* null mutant males. (B) Proportion of males that are fully infertile. Most *Dnaaf4* mutant males are infertile but this is rescued by the Dnaaf4-mVenus transgene (P=0.001, Fisher’s exact test) (C) Number of progeny per male, showing that rescued homozygous males are fully fertile compared with heterozygotes (P>0.9999, Kruskal-Wallis analysis followed by Dunn’s test for multiple comparisons). *n* = 10 males for each genotype. (C) Data for males in (B) plotted as number of progeny per male. A single Dnaaf4 homozygote gave progeny, perhaps due to being non-virgin at collection – 40 progeny compared with a mean of 96.9 for heterozygotes. (D) Fertility assay results showing a decrease in the number of fertile males in the *Dnaaf6* null mutant when compared to control groups (0.0001). *Dnaaf6* rescue did not produce progeny (P<0.0001) like that of the homozygous null mutants. *n* = 10 males per genotype. (E–I) Testes and associated male reproductive structures dissected from adult males and observed by light microscopy. Scale bars, 50 μm. (E) *Dnaaf4* heterozygote testis showing S-shaped motile sperm emerging from large (sperm-filled) seminal vesicle (black arrow). (F) *Dnaaf4* homozygote testis showing small (empty) seminal vesicle (black arrow) and absence of motile sperm. (G) Testis from *Dnaaf4* homozygote with *Dnaaf4*-mVenus transgene showing rescue of motile sperm production. (H) *Dnaaf6* heterozygote showing S-shaped motile sperm emerging from large (sperm-filled) seminal vesicle (black arrow). (I) *Dnaaf6* homozygote testes homozygote testis showing absence of motile sperm.

**Figure 6.**
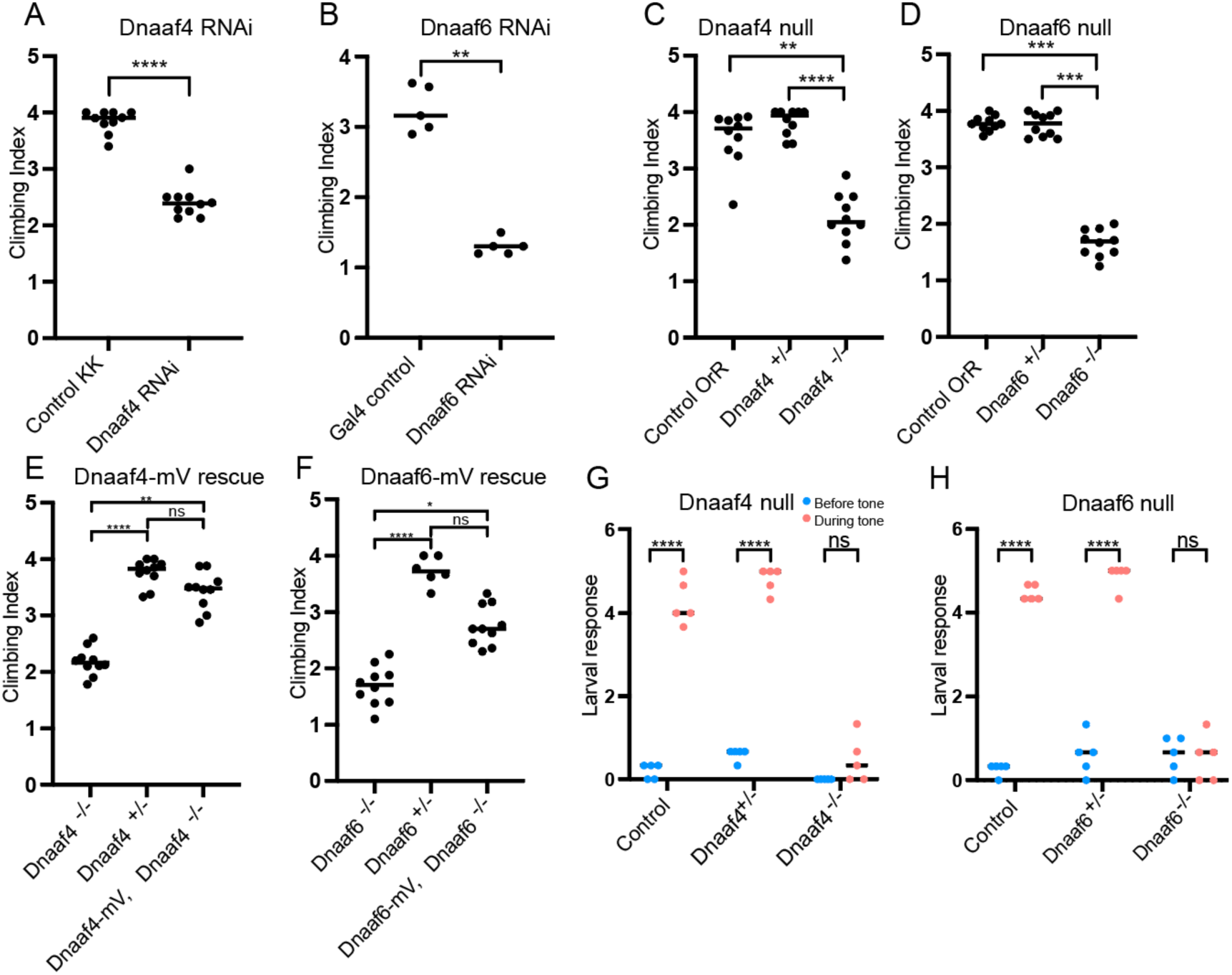
Knockdown and Null mutants of *Dnaaf4* and *Dnaaf6* have defective chordotonal sensory function. (A–F) Adult climbing assays for proprioceptive ability. Plots (with median and individual values), each point is a batch of 8-12 females, n=10 batches. (A,B) RNAi knockdown of *Dnaaf4* and *Dnaaf6* in sensory neurons (*scaGal4*) results in significant decrease in climbing ability. (C,D) Homozygote null adults for *Dnaaf4* and *Dnaaf6* have significantly decreased climbing ability compared with heterozygotes. (E,F) Rescue of null mutants. (E) *Dnaaf4*-mVenus transgene rescued the climbing ability of *Dnaaf4* null mutant flies, showing a significant increase in climbing performance when compared to null (P= 0.0012), restoring climbing ability to the same level as the heterozygotes (P=0.8130). (F) *Dnaaf6*-mVenus transgene partial restores climbing ability of *Dnaaf6* null mutants (P=0.0103), but not to levels seen in the heterozygote, although the latter difference does not reach significance (P=0.1282). (G,H) Plots (with individual and median values) showing hearing assay performances for *Dnaaf4*^-/-^ and *Dnaaf6*^-/-^ larvae in comparison to heterozygote and wild-type (OrR) controls. Number of larvae contracting before and during a 1000-Hz tone was measured. Individual points are batches of 5 larvae, n=5 batches. There is a significant difference between the number of larvae contracting before and during the tone (P < 0.0001) for control groups of both genotypes. There is no significant difference between the number of contractions occurring before and during the tone for *Dnaaf4* or *Dnaaf6* null mutants, indicating no behavioural response to stimulus. For climbing assays, significance was determined by Kruskal-Wallis followed by Dunn’s test for multiple comparisons. For hearing assay, significance was determined by two-way RM ANOVA and Sidak’s multiple comparisons test. Statistical significance on plots is indicated by asterisks: *, P≤0.05; **, P≤0.01; ***, P≤0.001; ****, P≤0.0001.

To confirm these phenotypes, CRISPR/Cas9 null mutants for *Dnaaf4* and *Dnaaf6* were generated, in which the open reading frame of each gene was replaced with the mini-*white* gene through homology-directed repair. For both *Dnaaf4* and *Dnaaf6*, homozygous null mutant flies are viable with no morphological defects, supporting the hypothesis that they are not required for general cellular functions. However, both *Dnaaf4* and *Dnaaf6* null males are infertile (Fig. 5B–D). Dissection of testes showed normal anatomy but a complete lack of motile sperm (Fig. 5E–I). In *Dnaaf4* null males, the development of motile sperm was rescued by the Dnaaf4-mVenus transgene (Fig. 5C,G). However, the Dnaaf6-mVenus transgene did not rescue the fertility of *Dnaaf6* males (Fig. 5D).

In a climbing assay, *Dnaaf4* and *Dnaaf6* homozygous null flies showed significant impairment compared to controls, consistent with defective chordotonal neuron function in proprioception (Fig. 6C,D). Climbing ability of null flies was restored fully or partially by Dnaaf4-mVenus and Dnaaf6-mVenus transgenes respectively (Fig. 6E,F).

To assess the auditory function of chordotonal neurons, a larval hearing assay was performed. Third-instar larvae normally respond to a 1000-Hz sine wave tone by momentarily contracting, a behaviour that requires functional dynein motors for mechanotransduction within chordotonal neuron cilia (zur Lage et al., 2021). Larvae homozygous for *Dnaaf4* or *Dnaaf6* mutations did not respond to a tone stimulus, consistent with functionally impaired chordotonal neurons in vibration sensing (Fig. 6G,H).

### Axonemal dyneins are defective in *Dnaaf4* and *Dnaaf6* mutant cilia

Overall, the phenotypes for *Dnaaf4* and *Dnaaf6* null flies are consistent with loss of dynein-driven motility in chordotonal neurons and sperm. To examine this further, TEM was performed on the chordotonal neuron array in the adult antenna (Johnston’s Organ) of *Dnaaf4* null mutant flies. This revealed largely normal neuronal structures including well-formed cilia, suggesting that there is no disruption of neuronal differentiation or general ciliogenesis. However, ODA and IDA were strongly reduced or absent (Fig. 7A,B). In antennae from *Dnaaf6* knockdown flies, TEM showed a strong reduction of IDAs and to a lesser extent ODAs (Fig. 7C,D).

**Figure 7.**
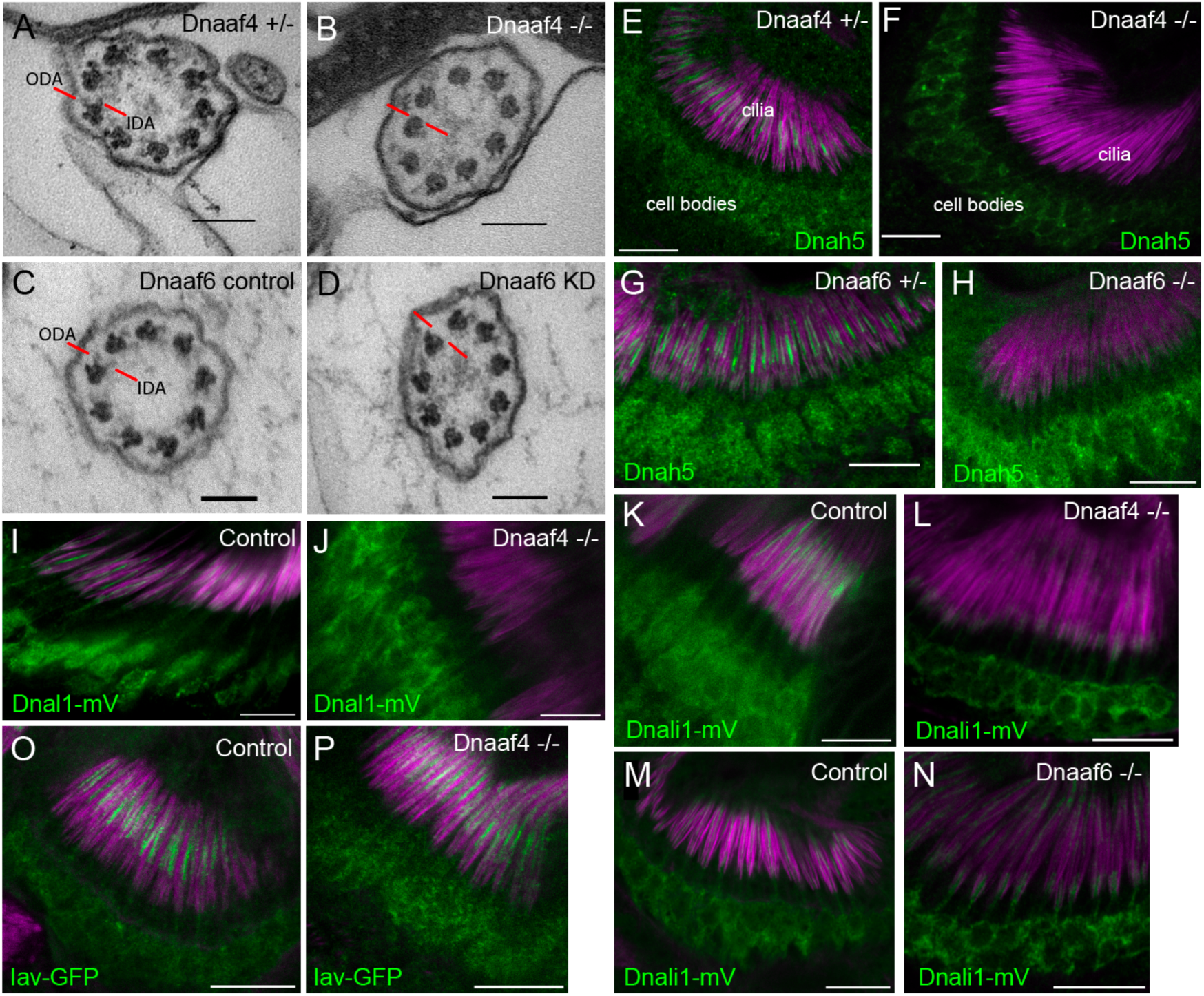
Defective dynein motor localisation in mutants. (A–D) TEM of chordotonal neurons in adult antennae, transverse sections of cilia showing 9+0 axonemal arrangement. (A) Control (*Dnaaf4*^+/-^ heterozygote) with ODAs and IDAs (red lines) on each microtubule doublet. (B) *Dnaaf4*^-/-^ homozygote showing severe loss of ODA and IDA structures from the microtubule doublets. (C) RNAi control (*scaGal4*, UAS-*Dcr2*, KK line) and (D) *Dnaaf6* knockdown (*scaGal4*, UAS-*Dcr2*, UAS-*Dnaaf6*RNAi). The latter shows a reduction of ODA and IDA. (E-P) Immunofluorescence of ODA/IDA markers (green) in differentiating chordotonal neurons of pupal antennae. All are counterstained with phalloidin, detecting the scolopale structures surrounding the cilia (magenta). (E–H) ODA heavy chain Dnah5 localisation in cilia is lost from *Dnaaf4*^-/-^ and *Dnaaf6*^-/-^ homozygote mutants (F,H) compared to controls (E,G), despite presence of protein in the cell bodies. (I,J) ODA marker, Dnal1-mVenus shows a similar loss of ciliary localisation in *Dnaaf4*^-/-^ homozygote (J) relative to control (I). (K–N) IDA marker, Dnali1-mVenus shows a partial loss of ciliary localisation in *Dnaaf4*^-/-^ and *Dnaaf6*^-/-^ homozygotes (L,N) relative to controls (K,M). (O,P) TRPV channel subunit Iav shows no difference in ciliary localisation between *Dnaaf4*^-/-^ homozygote (P) and control (O). Scale bars: (A–D) 100 nm, (E–P) 10 mm. Number of antennae imaged for IF: (E) n=7; (F) 7; (G) 6; (H) 5; (I) 5; (J) 10; (K) 5; (L) 9; (M) 8; (N) 9; (O) 6; (P) 7.

We extended these observations by examining the localisation of dynein markers in chordotonal neurons of pupal antennae. The ODA heavy chain, Dnah5, showed a complete loss of ciliary localisation in both *Dnaaf4* and *Dnaaf6* mutants (Fig. 7E–H). For *Dnaaf4*, similar loss was observed for the ODA light chain marker, Dnal1-mVenus (Fig. 7I,J). A marker of IDA subsets a,c,d, Dnali1-mVenus (light-intermediate chain 1), showed partial loss in ciliary localisation, which was more pronounced in *Dnaaf4* than *Dnaaf6* mutants (Fig. 7K–N). In contrast, the cilium localised TRPV channel subunit, Iav, was not altered in *Dnaaf4* mutants (Fig. 7O,P), suggesting that disruption of ciliary protein localisation is restricted to dynein complexes. Together, these observations suggest that both genes are required specifically for ciliary localisation of axonemal dyneins.

To investigate further, we assessed changes in protein abundance in *Dnaaf4* knock-down testes by label free quantitative mass spectrometry. In such experiments, a reduction in dynein chains has been considered consistent with instability resulting from defective cytoplasmic pre-assembly (zur Lage et al., 2018; zur Lage et al., 2021). Proteins detected in *Dnaaf4* knock-down testes were compared with control testes, and then filtered to concentrate on those associated with ciliary motility (dynein motors, nexin-dynein regulatory complex, radial spokes, etc (zur Lage et al., 2019)). As expected, Dnaaf4 protein is strongly depleted in knockdown testes ((log2(FC)=-8.69) (Fig. 8A). Of the other ciliary proteins detected, we found a small reduction in several ODA and IDA heavy chains, including kl-3 (orthologue: DNAH8, ODA), Dnah3 (DNAH3, IDA subsets a,b,c,e) and Dhc16F (DNAH6, IDA subset g). Also reduced were CG15128 (paralogue of TTC25, ODA docking complex), CG10750 (CCDC43B, MIA complex) and CG13168 (IQCD, Nexin-DRC). This may reflect a reduction in axonemal stability that appears to be characteristic of dynein loss in spermiogenesis (zur Lage et al., 2021). Interestingly, there is a small increase in Dnaaf2, which is one of the potential partners of Dnaaf4. To compare with the phenotype of another DNAAF, we also determined protein changes upon knockdown of TPR-containing *Spag1* (zur Lage et al., 2018). After filtering for motile ciliary proteins, we found very little difference in protein abundances between *Dnaaf4* and *Spag1* knockdown testes, suggesting that the roles of these DNAAFs are similar, or at least not distinguishable by this technique (Fig. 8B).

**Figure 8.**
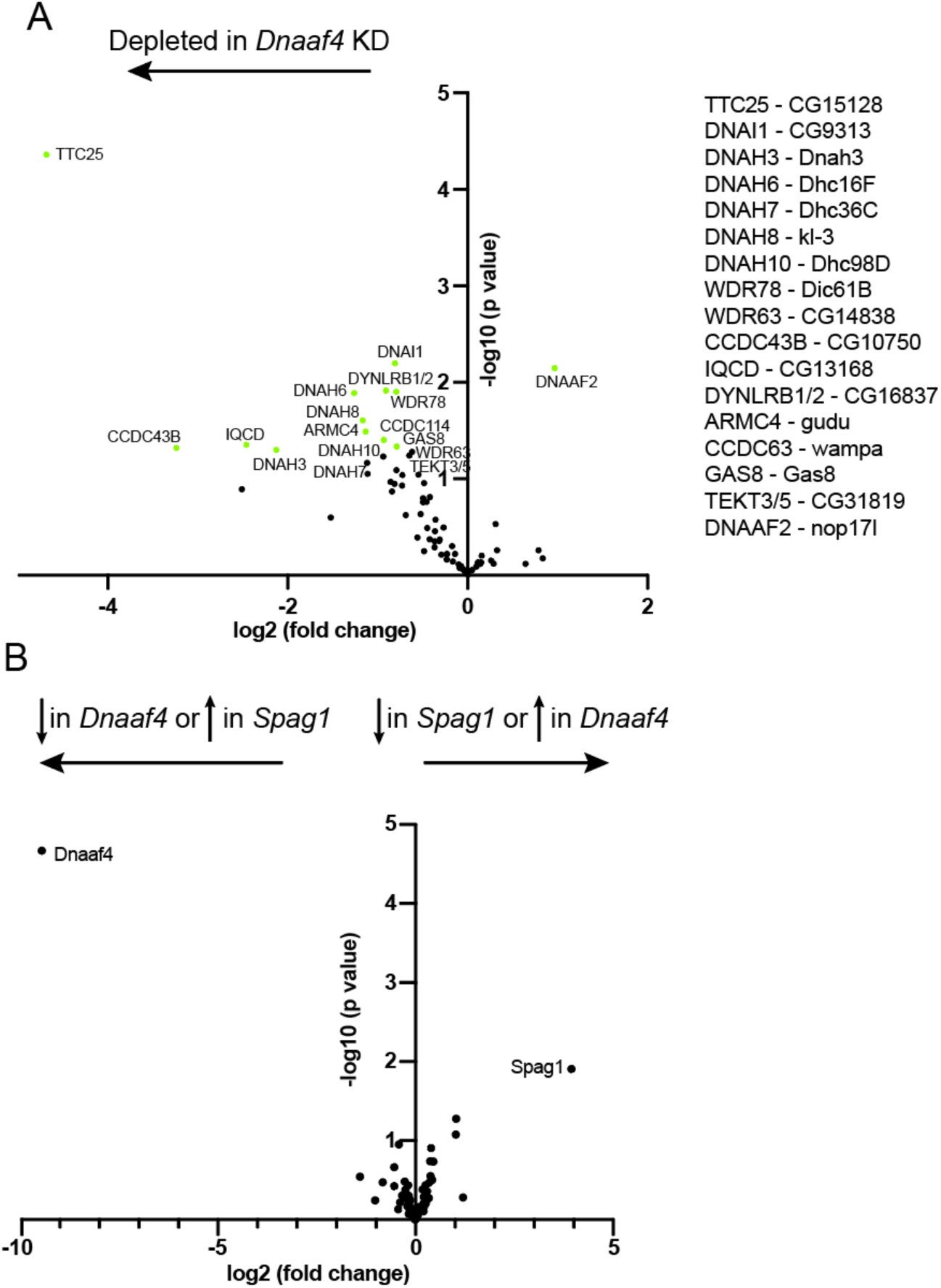
Proteomic changes in *Dnaaf4* mutant testes. (A) Volcano plot of motile cilia-associated proteins detected by MS in testes. To the left of the Y axis are proteins that are more less abundant in *Dnaaf4RNAi* KD (*BamGal4*, UAS*-Dnaaf4RNAi*) testes compared with *BamGal4* control (depleted); to the right are proteins that are more abundant than in the control. Dnaaf4 protein itself is strongly depleted as expected (log2(FC)=-8.69, -log10(p value)= 4.39) but for clarity it is not shown on plot. Proteins with -log10(p value)>1.3 (green points) are labelled with names of human homologues. The *Drosophila* gene names are shown to the right. *n* = 30 pairs of testes/replicate; 4 replicates per genotype. (B) Volcano plot comparing motile cilia-associated proteins detected in testes from *Dnaaf4* knockdown testes compared with *Spag1* knockdown testes (*BamGal4*, UAS*-Spag1RNAi*). The only proteins showing significant difference in abundance are Dnaaf4 and Spag1 themselves. Significance was determined using the Empirical Bayes method. *n* = 30 pairs of testes/replicate; 4 replicates per genotype.

## DISCUSSION

*Drosophila Dnaaf4* and *Dnaaf6* are both required for axonemal dynein localisation within cilia, showing that despite the truncated nature of *Dnaaf4*, there is conservation of the roles assigned to homologues in other organisms. Physical evidence supports the possibility that they perform this role as part of an R2TP-like complex in *Drosophila*. On the other hand, for neither gene do we find evidence of function beyond the differentiation of motile cilia, suggesting that in *Drosophila* at least, the role of these genes is specific to axonemal dynein assembly.

Vertebrate DNAAF4 is predicted to recruit HSP90 via its TPR domain, and we show that mouse Dnaaf4 is able to bind Hsp90. It is remarkable, therefore, that despite apparent conservation of function as an Hsp90 co-chaperone, *Drosophila* Dnaaf4 protein lacks the TPR domain. Perhaps an accessory TPR-containing protein works with *Drosophila* Dnaaf4. Interestingly, *Drosophila* Spag1 is also strongly truncated, but in this case the truncation retains the TPR domain and not much else (zur Lage et al., 2019). Does Spag1 work in partnership with Dnaaf4? Our proteomic analysis of knockdown testes suggests that Dnaaf4 and Spag1 have similar phenotypes. However, affinity purification analysis did not detect Spag1 as a Dnaaf4-interacting protein. On the other hand, this analysis also did not detect interaction with Dnaaf6, and so the conditions of the assay may not be conducive to identifying Dnaaf4 protein interactors efficiently.

There are questions regarding the role of the DNAAF4 TPR domain humans too, since the protein exists in several isoforms with varying numbers of repeats in its TPR domain (Fig. 1B). While isoform-a (which associates with DNAAF2) has a full 3-repeat TPR domain that is likely to be essential for HSP90 binding (Maurizy et al., 2018; Takar et al., 2013), isoform-c (which associates with DNAAF6) has only a single repeat (Maurizy et al., 2018; Paff et al., 2017). It seems unlikely that the limited TPR domain of isoform-c can bind HSP90 directly, and so it may not differ functionally from the *Drosophila* protein so strongly after all.

Interestingly, *Drosophila* truncated Dnaaf4 resembles the protein that would potentially be synthesised from the human gene bearing the pathogenic mutation detected in PCD: in the original report, 7 out of 9 *DNAAF4* variants in PCD patients were nonsense mutations predicted to encode a truncated protein lacking TPR domains (Tarkar et al., 2013). However, as nonsense-mediated decay (NMD) of the transcript is thought to occur, it is likely that no protein is produced. The finding that *Drosophila* truncated *Dnaaf4* is functional without a TPR domain raises the possibility that inhibition of NMD could restore some function to PCD patients with DNAAF truncating mutations, even if the protein produced lacks the TPR domain. On the other hand, we found in our heterologous expression system that the full TPR domain of mouse Dnaaf4 was required for strong interaction with Dnaaf6.

We find that *Drosophila* mutants of *Dnaaf4* and *Dnaaf6* show similar loss of dynein markers. While the markers available in *Drosophila* are limited, this finding supports them working in the same complex. There is a strong loss of ODA markers (Dnal1 and Dnah5 homologues) but a partial loss of IDA marker, Dnali1. This chain is predicted to be a subunit of single-headed IDA subsets a, c and d, although it is not certain that d exists in *Drosophila* (zur Lage et al., 2019). In comparison, electron tomography analysis of human PIH1D3-mutant respiratory cilia showed a loss of subset g but no effect on subsets a or c (Olcese et al., 2017).

Mutations of the Dnaaf4 homologue in *Chlamydomonas* resulted in strong reduction of most IDA subsets but a weak reduction of subset a (Yamamoto et al., 2017). In other organisms, homologues of these DNAAFs have also been proposed to have a role in the assembly of subset g. For further precision on the subsets affected in *Drosophila*, it would be desirable to generate heavy chain markers for IDA subsets such as antibody raised against the IDA heavy chain DNAH6 homologue, *Dhc16*. Dnaaf4 is proposed to function with Dnaaf2 in addition to Dnaaf6, and it is not known whether this would be responsible for the assembly of other dynein complexes. Given that Dnali1 expression appears lower in the *Dnaaf4* mutant than the *Dnaaf6* mutant, this may also suggest a role for Dnaaf4 partners with proteins in addition to Dnaaf6.

Several DNAAFs are suspected of having additional non-ciliary functions. For example, mice *DNAAF2* homozygotes are reported to be embryonic lethal (Cheong et al., 2019) consistent with wider roles, and it may be significant that only a small number of PCD patients have been identified with mutations in *DNAAF2* (Omran et al., 2008). In *Drosophila, Dnaaf2* (*nop17l*) appears to be widely expressed in embryos (zur Lage et al., 2019), supporting the possibility of widespread roles for this *DNAAF*. In contrast, *Drosophila Dnaaf4* is specifically expressed in motile ciliated cells supporting the hypothesis that has no other roles than facilitating axonemal dynein assembly. In this light it is interesting to consider the roles proposed for vertebrate *DNAAF4*. Truncating mutations of *DNAAF4* were first identified as a candidate causative gene for dyslexia through a role in brain development and maturation (Taipale et al., 2003). Based on rodent models, it has been proposed that *DNAAF4* mutation affects neuronal migration in the developing neocortex (Wang et al., 2006) The link between *DNAAF4* and dyslexia requires further confirmation since this gene did not associate with dyslexia in follow-up studies on other populations (Marino et al., 2005; Scerri, 2004). It is not immediately clear how such a phenotype depends on ciliary motility, raising the possibility that *DNAAF4* may have additional non-ciliary roles. Alternatively, a potential role in neuronal migration/dyslexia could also be an indirect effect of a motile cilia defect, since ciliary motility is required for CSF flow (Kumar et al., 2021).

Another intriguing possibility arises from the observation that neuropsychiatric disorders such as schizophrenia, autism and dyslexia have been connected to left-right asymmetry (Trulioff et al., 2017; Valente et al., 2014), which is determined via motile cilia in the embryonic node. Indeed, a recent case report of mutations in the dynein heavy chain genes, *DNAH5* and *DNAH11* has raised the possibility of a link between situs inversus and developmental dyslexia (Bieder et al., 2020).

## Author Contributions

JL conducted many of the experiments and data analysis, contributed to experimental design; PzL conducted S2 cell analysis and contributed to mVenus construction and analysis; AvK provided the mass spectrometry analyses; APJ conducted data analysis, contributed to experimental design, and wrote the manuscript. All authors contributed to manuscript revision, read and approved the submitted manuscript.

## Acknowledgments

We thank IfeOluwa Taiwo and Panagiota Stefannopoulou for preliminary contributions, and Tracy Davey of Electron Microscopy Research Services, Newcastle University Medical School for TEM. Stocks obtained from the Bloomington *Drosophila* Stock Center were used in this study.

## Data Availability Statement

The mass spectrometry datasets generated for this study can be found in the ProteomeXchange Consortium via the PRIDE (Perez-Riverol et al., 2019) partner repository with the dataset identifier PXD033608.

